# RATTACA: Genetic predictions in Heterogeneous Stock rats offer a new tool for genetic correlation and experimental design

**DOI:** 10.1101/2023.09.18.558279

**Authors:** Benjamin B. Johnson, Thiago M. Sanches, Mika H. Okamoto, Khai-Min Nguyen, Clara A. Ortez, Oksana Polesskaya, Abraham A. Palmer

## Abstract

Genetic correlations between traits are a common first step in studies identifying causal genetic pathways and mechanisms. Using this framework with inbred or selected lines, however, requires intensive labor investment through breeding and phenotyping, and is prone to confounding, as observed trait correlations do not necessarily reflect a causative genetic architecture shared between distinct populations. When drawn from a single outbred population, genetic trait predictions offer a viable alternative to experimental phenotyping and can be used to identify putative genetic correlations when samples with divergent trait predictions also diverge in a second measured trait. Here, we present a novel research paradigm and service called RATTACA, in which genotypes from Heterogenous Stock (HS) rats are used to predict trait values using linear mixed models. These predictions are used to select samples of individuals with high and low extreme trait values, facilitating (1) *a priori* sampling of desired trait values without oversampling across phenotypic space and (2) easy identification of putative genetic correlations between predicted and newly measured traits. We validated prediction models using four example phenotypes with measured trait values and found sufficient accuracy to distinguish extreme trait samples, even when using a small number of genome-wide variants (n = 50,000) for traits with modest heritability (*h^2^* = 0.13). Given genotypes and trait measurements available through previous research in HS rats, we propose RATTACA as a service to reliably predict more than 80 behavioral and physiological traits.

## Introduction

Genetic correlations between traits are a foundational tool used to help identify causal mechanisms linking genotype and phenotype (Falconer 1960). However, genetic correlations can be difficult to effectively implement due to (1) confounding effects introduced by study design, (2) resource and labor-intensive laboratory practices required for these studies, and (3) the large sample sizes needed to produce correlations with adequate statistical power. Simple (observed) correlations between traits do not necessarily reflect a shared genetic architecture, and thus do not inherently suggest a genetic correlation. As such, observed correlations risk confounding genetic and environmental effects on phenotype (Falconer 1960). For example, in selected lines, traits are selected to diverge between strains, but correlations with a second, unselected trait observed across strains need not reflect a common genetic basis for both traits. Rather, these correlations may result from drift in the unselected trait between separate populations. Comparing traits between larger panels of multiple inbred strains can help disentangle the contribution of genetics to phenotypic variation (Silver 1995). However, such solutions require intensive breeding programs that cost substantially in effort, time, resources, and funds (Heffner et al. 2010).

Like breeding, phenotyping is a similarly labor-intensive pursuit. Phenotyping often requires substantial effort by highly trained scientists invested into observational experiments or, alternatively, automated technologies (e.g., video recordings, sensors, etc.) that are complicated and expensive (Whishaw and Kolb 2004). To make statistical comparisons between extreme trait values, studies have traditionally required panels of inbred strains or selectively bred lines. Both of these approaches are difficult or expensive to implement. Carefully maintained outbred populations can reduce effort to one line and eliminate the confounding environmental effects that arise between separate lines, but phenotyping outbred samples requires larger sample sizes to capture adequate phenotypic variation necessary for robust genetic correlations (M. F. Festing 1976; M. F. W. Festing 2014). Given the substantial methodological constraints that underlie any attempt to link genotype to phenotypes, these research programs are limited to investigators with access to considerable resources. Clearly, if we can minimize the obstacles to both effective phenotyping and genetic correlation, future studies will have greater power to uncover the genetic architecture of a diversity of biological traits.

Continuing advances in genomic sequencing technologies have made high-throughput genotyping, which can be used to impute millions of genome-wide single nucleotide polymorphism (SNP) genotypes, efficient and economical. This opens a novel avenue for experimental design in quantitative genetics: genetic trait prediction (Jeon et al. 2023). Genetic trait prediction is an established practice in agricultural and human genetics that has found numerous applications such as predicting crop yields and human disease risk (Bhat et al. 2016; Ma and Zhou 2021). In one common implementation of this practice, correlations between sample genotypes and phenotypes are used in linear mixed models to estimate the best linear unbiased predictor of per-variant genetic effect sizes (G-BLUP) (Fernando and Grossman 1989). These estimates can in turn be used to predict per-individual breeding values - the total genetic contribution to the phenotype - for a given trait of interest (Heffner, Sorrells, and Jannink 2009). When produced using an appropriate sample and model, these predictions can reliably identify individuals with extreme phenotypic values. Identifying these individuals without having to measure the trait can thus enable researchers to select an *a priori* sample of desired genotypes to experimentally phenotype, minimizing experimental effort and cost by eliminating the need for oversampling or inbreeding in search of desired phenotypic values.

The value of a given predictive model invariably depends upon its accuracy. This accuracy will stem primarily from SNPs that are either (1) in linkage disequilibrium (LD) with causal loci or (2) reflective of the relationship structure between the training sample and the sample to be predicted (Habier, Fernando, and Dekkers 2007). While genome-wide association studies (GWASs) have identified numerous SNPs associated with complex traits, these methods are not good for capturing causal loci of small effect (Holland et al. 2020; Uffelmann et al. 2021). As such, significant variants identified by GWASs account for only a fraction of the total genetic variation underlying a given trait (Manolio et al. 2009), and models fit on these SNPs tend to predict trait values poorly (de Los Campos et al. 2013). In contrast, genome-wide SNP samples can explain a greater proportion of phenotypic variance by capturing the joint contribution of many variants of small effect (Yang et al. 2010). Extremely large SNP samples, however, can make predictions computationally difficult, and may not improve prediction accuracy when few SNPs are in LD with causal variants (Ober et al. 2012). Optimized models thus require identifying a sufficient subset of genomic variants of small to moderate effect to provide reliably accurate predictions (Bermingham et al. 2015). While the performance of predictions is ultimately constrained by the genetic architecture (e.g., the SNP composition and heritability) of a given phenotype, model performance for a given trait can be maximized by identifying this sufficient set of variants that reliably predicts well across novel samples, all while balancing computational demands.

Here, we introduce a novel application of genetic predictions to identify genetic correlations and inform the sampling of extreme traits in an outbred laboratory population of rats (*Rattus norvegicus*). This population, the N/NIH Heterogeneous Stock (HS) (Hansen and Spuhler 1984), is a longstanding outbred strain regularly used in GWASs of behavioral and physiological traits (Solberg-Woods and Palmer 2019; Chitre et al. 2020); (Polesskaya et al., n.d.). By leveraging the extensive genetic and phenotypic data accrued through this population we have developed a novel research service, termed RATTACA (**<u>RAT T</u>**rait **<u>A</u>**scertainment using **<u>C</u>**ommon **<u>A</u>**lleles). RATTACA comprises a concerted breeding program that provides regular, generational genetic prediction on naive rats to facilitate researchers’ *a priori* sampling of genotypes with desired expected trait values. Here, we outline our workflow for producing rats and trait predictions, which we validated and optimized by testing model predictions on samples with known values for four example phenotypes. Using this framework, we can offer reliable and computationally tractable genetic predictions for dozens of traits measured across multiple behavioral paradigms, as well as physiological and -omics traits. By demonstrating the efficacy of predictions across a breadth of potential traits, we present RATTACA as a valuable new resource for experimentalists pursuing the genetic architecture of behavioral phenotypes using HS rats as a model, and as a novel framework for sampling design in other experimental systems linking genotype and phenotype.

## Methods

### HS rats colony and breeding program

To provide regular generations of naive rats to researchers, we established a new colony of Heterogeneous Stock rats at the University of California San Diego. Operating as part of the P50 Center for Genetic Studies in Outbred Rats (Polesskaya et al., n.d.), the UCSD Heterogeneous Stock colony (termed ‘HS West’; strain ID McwiWfsmAap:HS #155269102, RID:RGD_155269102) was initiated in July 2022 using 64 breeder pairs from the HS colony maintained at Wake Forest University (colony ID NMcwiWFsm #13673907, RRID:RGD_13673907) (Sanches et al., n.d.). The HS West colony operates for the purpose of providing HS rats for genetic prediction and other experimental uses. As of August 2023, the colony is in its 99th generation since the inception of HS rats strain in 1984. Generations are produced on a three month cycle, offering four batches per year of about 500 rats per batch for regular selection into experiments by collaborators.

Each generation cycle is characterized by three primary milestones: (1) pairing breeders, (2) weaning litters, and (3) assigning animals to projects. We begin each cycle by pairing 70 breeding pairs over 10 days. Litters are born approximately three weeks after pairing, at which point we remove and euthanize the male breeder and collect genetic material (tail and spleen) for future genotyping. Weaning occurs at three weeks of age, at which point pups are separated from the mother. We keep a minimum of four animals of each sex per litter, each of which is ear punched for in-cage identification and subcutaneously injected with a unique identifier RFID chip. Critically, we collect the ear-punched tissue for immediate genotyping. After weaning, the female breeder is also euthanized, with genetic material collected for genotyping.

Prior to genotyping, investigators can submit a request for a desired number of rats with extreme values for a specific trait. We then use the current generation’s genotypes to perform genetic prediction on all individuals for all requested traits. Rats are then assigned to respective investigators based on predicted trait values using a custom algorithm coded in Python. Briefly, we aim to maximize sample differences between high and low trait values assigned to a given project, while avoiding biasing samples by sex and while minimizing the number of siblings. To do so, we first identify each individual’s rank order of predicted trait value for the desired trait, then select the set of available rats with extreme high and low ranks to produce two samples expected to differ significantly in mean trait values (Fig. 1). This set of available rats will depend on a project’s requested sample size and on whether individuals have already been provided to another project. To minimize sample biases, we provide a maximum of one male and one female from each litter and provide approximately equal proportions of males and females in either sample, unless the request specifies otherwise.

**Figure 1.**
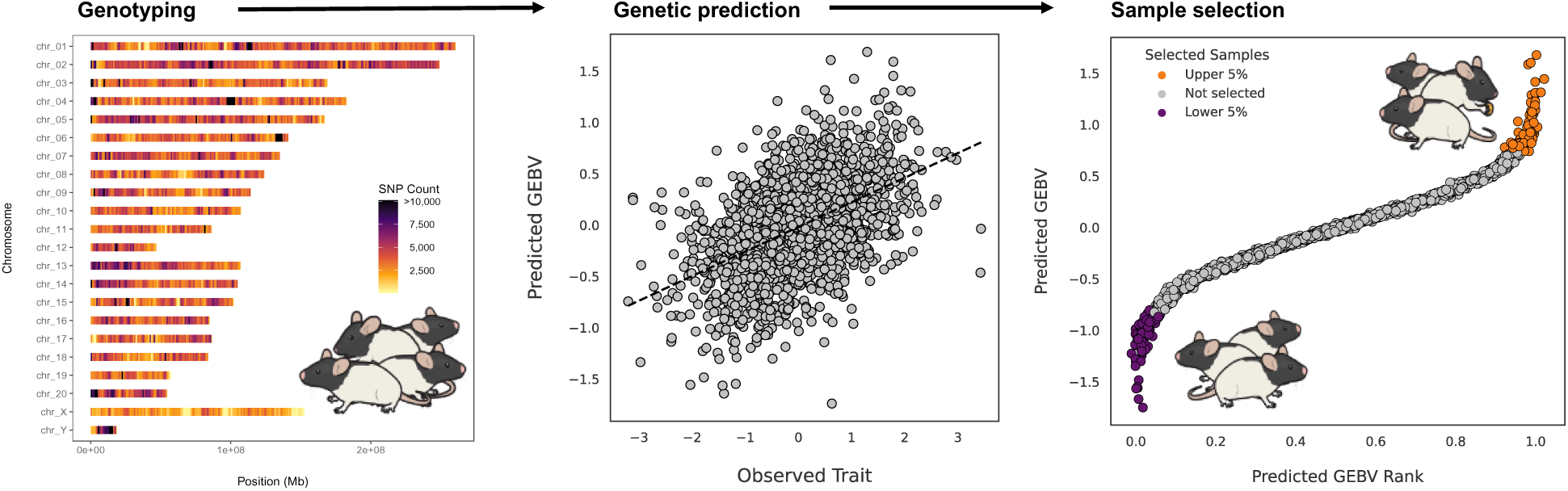
The RATTACA paradigm for genetic prediction and sample design. With each successive generation of HS rats produced at the HS West colony, weaned pups are genotyped at >7M genome-wide SNP variants. For a given trait of interest, all genotypes and phenotypes previously measured in HS rats are used to train a predictive G-BLUP model, which is then fit using the population of new genotypes for genetic prediction. Predicted genomic-estimated breeding values (GEBVs) are ranked-ordered across the population, and samples are drawn from extreme GEBVs (here, the highest and lowest 5% of the distribution) to produce experimental samples expected to differ in mean trait value.

Aside from rats provided to investigators, we invariably reserve one male and female from each litter for subsequent breeding. All retained breeders have known genotypes and pedigrees spanning ∼45 generations. With this information, we use the R package ‘breedail’ (Carbonetto 2014) to assign new breeder pairs with the lowest shared kinship coefficient. Following assignment, breeders are paired to initiate a new cycle of breeding, weaning, and prediction.

### Genomic data preparation

All RATTACA genotypes are produced following a common protocol (Supplemental Material). Briefly, we extract genomic DNA from earpunch tissue using the Agencourt DNAdvance Kit (Beckman Coulter Life Sciences, Indianapolis, IN), measure DNA concentrations and assessed extraction quality using a NanoDrop 8000 spectrophotometer (Thermo Fisher Scientific, Waltham, MA), and use concentrations to normalize sample quantity prior to library preparation. Genomic libraries are prepared using the Twist 96-Plex Library Preparation Kit (Twist Bioscience, San Francisco, CA). We quantify library concentrations using a Qubit 3.0 Fluorometer (Thermo Fisher Scientific, Waltham, MA) and assess average fragment sizes using the Agilent 4200 TapeStation (Agilent Technologies, Santa Clara, CA). Following quantification, we normalize all libraries to 10mM and pool all for sequencing using paired-end 2x150 bp reads over one lane of an S4 flowcell on an Illumina NovaSeq 6000 DNA sequencer (Illumina, San Diego, CA) run at the UCSD Institute for Genomic Medicine Genomics Center. Finally, we demultiplex sequenced libraries and imputed sample genotypes following Chen et al. (Chen et al., n.d.) to produce a common set of SNP genotypes for all samples.

While RATTACA predictions are an ongoing endeavor, all model optimizations and trait predictions presented in this manuscript were conducted using available genotypes and trait measurements previously produced for various GWASs at the Center for Genetic Studies in Outbred Rats (www.ratgenes.org; Polesskaya et al., n.d.). These sample genotypes were all produced following the same analytical pipeline as above (Chen et al. 2023).

### Phenotypes

We trained and validated G-BLUP predictions on four example phenotypes (Table 1) using trait measurements and genotype data produced in prior experiments. All phenotypic data used for these analyses are experimental measurements produced as part of the Center for Genetic Studies in Outbred Rats (Polesskaya et al., n.d.), whose primary focus is on addiction-related behaviors underlying substance use disorder (SUD). We selected these traits for three reasons: (1) They span a diversity of psychological and physiological categories of potential interest to investigators studying the genetics of SUD; (2) The narrow-sense heritability of these traits (*h^2^*, calculated during previous GWAS analyses) varies broadly, allowing us to test the effect of *h^2^* on genetic prediction accuracy; and (3) The large sample sizes available for each trait allows us to test the effect of training sample size on prediction accuracy.

**Table 1.** Four example traits used for model training and validation. Narrow-sense heritability (*h^2^*) spans from modest (NRR) to high (Mass) in this set, allowing tests for genetic prediction accuracy across a wide range of scenarios. Trait categories are also highly variable, demonstrating the general utility of RATTACA to a diversity of potential phenotypes.

| Trait | Acronym | Category | N | $h^2$ | Description | Citation |
| --- | --- | --- | --- | --- | --- | --- |
| **Body mass** | Mass | Physiology | 5670 | 0.467 | Body mass measured at the time of sacrifice at approximately 200 days of age | Chitre et al. 2020 |
| **Distance traveled in a novel environment** | DNE | Behavior: novelty seeking, exploratory behavior | 2509 | 0.274 | Total distance traveled during 18 minutes in standard plastic cage (42.5×22.5×19.25 cm) | Garcanz et al. 2012 |
| **Visual stimulus-reinforced responding** | VSSR | Behavior: sensation seeking | 2405 | 0.282 | Active snout-poke responses for 5-sec light onset in the last three 18-minute sessions during the light reinforcement phase | Garcanz et al. 2012 |
| **Non-reinforced responding** | NRR | Behavior: impulsivity, habituation | 2406 | 0.129 | Inactive snout-poke responses with no consequences in the last three 18-minute sessions during the pre-exposure (with no light reinforcer) | Garcanz et al. 2012 |

We conducted predictions on three trait categories: body weight, locomotor activity, and light reinforcement. We used the dataset of Chitre et al. (2020) to analyze body weight (mass), a commonly measured trait that covaries with a wide array of physiological phenotypes (Chitre et al. 2020). Locomotor activity measurements were conducted as described by (Gancarz, Robble, et al. 2012). We analyzed distance traveled in a novel environment (DNE), an exploratory behavior reflective of sensation seeking and reactions to novelty in rodents (Gancarz, Robble, et al. 2012). Light Reinforcement phenotypes were measured following (Gancarz, Ashrafioun, et al. 2012). These traits underlie behavioral regulation related to drug abuse susceptibility, including attention, impulsivity, habituation, and sensitization (Gancarz, Ashrafioun, et al. 2012; Wang et al. 2020; Gancarz, Robble, et al. 2012). We analyzed visual stimulus-reinforced responses (VSSR) and non-reinforced responding (NRR) from the dataset of Gancarz et al. (2012). All protocols were approved by each project’s respective Institutional Animal Care and Use Committees.

### Trait predictions

For all trait predictions, we used the ‘mixed.solve()’ function in the R package ‘rrBLUP’ v4.6.2 (Endelman 2011), a program commonly implemented in genomic selection in crop sciences. Briefly, this method predicts complex traits using marker data as random effects in linear mixed models. To do so, this marker-based (G-BLUP) method calculates BLUP solutions for individual SNP effects on a given sampled phenotype, whose sum produces an individual genotype’s genomic-estimated breeding value (GEBV). For all predictions, we followed a general framework of fitting models to a training set of measured genotype and phenotype data, then employing the fitted model to predict GEBVs using genotypes in a test sample (see genomic sampling details below). We assessed model performance by comparing predicted GEBVs to known phenotype values in the test sample. We defined model performance as prediction accuracy, specifically defined as the Pearson correlation (*r*) between observed test phenotypes and GEBVs predicted by the model, with higher correlations reflective of greater accuracy and superior model performance.

All predictions were performed on standardized residuals derived from linear models fit to each trait sample. Prior to model predictions, we ensured normality in all trait samples by quantile normalizing each trait’s measured values to a Gaussian distribution. We identified all potential experimental covariates (including sex, age, generational cohort, and cage) and fit linear regressions between normalized trait values and each covariate. We retained all covariates with *r^2^* ≥ 0.02 to include in a ‘final’ model of important covariates. We then fit this final model on the sample of normalized trait values and retained the model residuals. Finally, we quantile normalized these residuals to use as our per-individual empirical trait observations in subsequent analyses.

### Genome sampling

Regular genetic prediction of multiple traits using a dense sample of genome-wide SNP variants has the potential to be computationally intensive. Further, because most variants are non-independent due to linkage disequilibrium (LD), genome-wide SNP samples may overrepresent genomic regions with higher LD, resulting in biased predictions no longer representative of genome-wide ancestry. As such, an LD-informed approach to subsampling the genome may be best to optimize genetic predictions. To this end, we sought to determine if accurate, reliable predictions can be produced using computationally manageable subsets of genomic variants by sampling genome-wide SNPs both randomly and by removing redundant variants in LD (‘LD pruning’).

To characterize the effect of SNP sample size on accuracy, we extracted random samples of 10, 100, 1,000, 10,000, 50,000, 100,000, 300,000, and 1,000,000 SNPs from sample genotypes using PLINK v1.9 (Chang et al. 2015). For each trait and genomic sample size, we tested model predictions on 10 independent random samples of rats. Each sample included 2,300 rats, 60% of which (1,380 individuals) were used for model training and 40% (920) for model testing. For LD pruning, we used the ‘--indep-pairwise’ option in PLINK. This method prunes SNPs from a sliding window of variants by checking the correlation between all variant pairs, removing one variant from each pair whose squared correlation (*r^2^*) exceeds a given threshold value. To determine optimal parameters for LD pruning, we compared predictions using 12 parameter combinations. We sampled the genome using r^2^ thresholds of 0.5, 0.99, and 0.9999, and for each threshold we used window sizes of 50, 100, 1,000, and 10,000 SNPs, and a sliding window of 1/10 the window size. For each parameter combination, we pruned SNPs and tested model predictions on 50 independent random samples of 2,300 rats, split into training and test sets as above. In each case, SNPs were pruned solely from the training set to ensure each replicate represented an independent SNP sample on which to test model predictions. Because extremely rare variants with a low minor allele frequency (MAF) are uncommon in HS rats (Munro et al. 2022); (Sanches et al., n.d.), all LD pruning and random sampling was conducted on the entire set of SNP genotypes per individual, without prior filters to exclude low-MAF SNPs.

To identify an optimal method for subsampling genotypes, we compared models trained on LD-pruned genotypes to those trained on random SNP samples. Because prediction performance was statistically equivalent between all LD pruning parameters (see Results), we proceeded with comparisons using only an r^2^ threshold of 0.99, a window size of 1,000 SNPs, and a sliding distance of 100 SNPs. For each trait, we randomly sampled the same number of variants as retained following LD pruning. We repeated sampling and model testing on 50 independent random samples of both rats and SNPs as above. To test for differences in model performance between genomic sampling strategies, we fit a linear model testing model performance as a function of trait plus sampling method. From this model, for each trait we compared estimated marginal means for either sampling method using a Tukey adjustment for multiple pairwise comparisons, implemented with the R package ‘emmeans’ v1.8.8 (Lenth 2023).

### Population sampling

To evaluate the effect of changing population sample sizes on prediction accuracy, we fit models to a range of training samples using a common SNP sample size. For each trait, we fit models on training samples of 10 rats, 50-450 rats in increments of 50, and 500-2,000 rats in increments of 100 (26 training sizes total). For each training sample we conducted 15 replicates using independent random samples of 50,000 SNPs. In all cases, we fit trained models onto a test sample of 300 rats to test prediction accuracy as a function of training sample size.

### Trait selection

Under its standard operation, RATTACA employs the entire available sample of individuals genotyped and phenotyped for a given trait in order to train predictions on naive, unmeasured samples. To simulate the process of selecting extreme trait predictions for inclusion in experiments, we predicted each of our four traits using the entire available sample of phenotyped rats, split into 70% training and 30% test sets. We identified the rank order of each predicted GEBV and ‘selected’ samples in the top 5% and bottom 5% of ordered values for inclusion in a hypothetical project (Fig. 1). For each trait, we tested our ability to distinguish between extreme samples by comparing the true, observed trait values in either selected sample using a student’s T-test and Bonferroni correction for four traits.

## Results

G-BLUP models predicted GEBVs with sufficient accuracy to distinguish between extreme samples (upper and lower 5%) for all tested traits (Fig. 2). While our selected extreme samples regularly overlap in their distribution of observed trait values (Fig. 2A), the mean of these samples differed significantly in all cases (*P* = 1.09 x 10^-4^ −4.87 x 10^-35^) (Fig. 2B). Using a common SNP sample size, model prediction accuracy increased with both training sample size and trait heritability (Fig. 3A). Accuracy was also positively correlated with the number of SNPs sampled (for a common training population size), however this relationship plateaued for samples greater than ∼10,000 SNPs (Fig. 3B). We found no difference in prediction accuracy between models fit on LD-pruned or randomly sampled SNP sets (Fig. 3C), as both methods were equivalent for all traits (*P* = 0.352; Supplementary Material).

**Figure 2.**
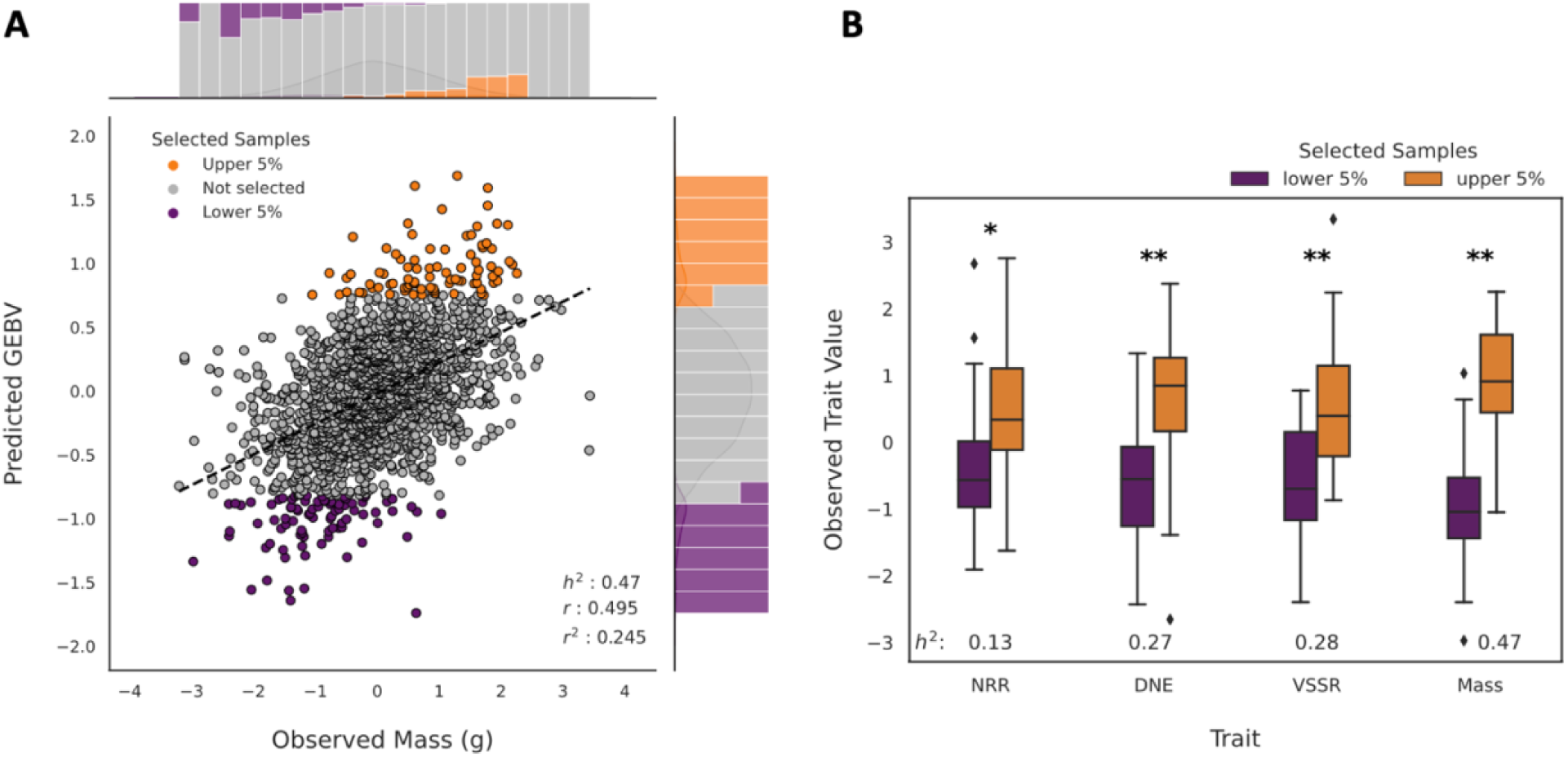
Trait predictions successfully distinguish extreme samples with distinct mean trait values. G-BLUP predictions of genomic-estimated breeding values (GEBVs) correlate with experimentally observed phenotypes (**A**), here using body mass as an example. Selecting the upper and lower 5% of predicted GEBVs by definition produces non-overlapping samples of GEBVs (right marginal histogram). Because predictions are imperfect (non 1:1 correlations), selected samples overlap in their distribution of trait values (top marginal histogram), but prediction accuracy is high enough that mean values are statistically distinct (**B**). The difference between the high and low groups increases with increasing trait heritability (*h^2^*). All differences between observed sample means are statistically significant (t-test; single asterisk *P* < 1×10^-3^, double asterisks *P* < 1×10^-6^).

**Figure 3.**
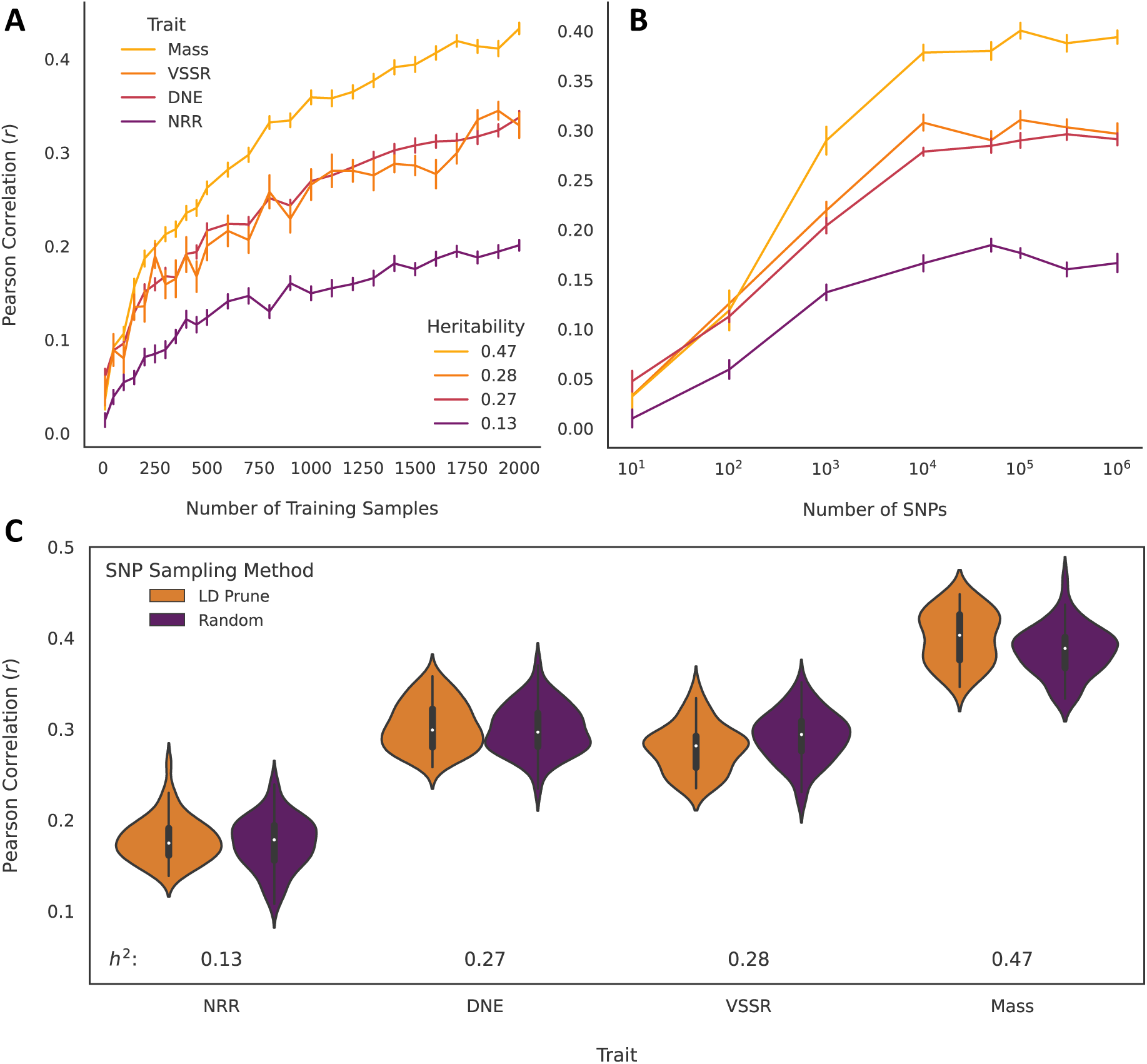
Population and genomic sample size positively influence trait prediction accuracy, but genomic sampling method does not. Prediction accuracy (the Pearson correlation between experimental measurements and predicted GEBVs) increases for all traits as training sample size used to fit the prediction model increases (**A**). Accuracy also increases when increasing the number of SNPs used for prediction, but only at lower SNP samples (< ∼10,000 SNPs), above which accuracy remains generally consistent (**B**). Accuracy is not influenced by variant sampling method, as performance is equivalent between models fit to SNP samples produced either randomly or by LD pruning (**C**). Pairwise *P* < 0.05 for all traits following a Tukey correction for multiple comparisons. Error bars in B and C are the standard error of the mean prediction accuracy.

## Discussion

### Genetic prediction as a novel approach to experimental design

Genetic correlations between traits are frequently employed in model systems as a first step towards identifying causal genetic pathways and mechanisms. Using this framework with inbred or selected lines, however, requires intensive labor investment through breeding and is prone to confounding, as observed trait correlations do not necessarily reflect a causative genetic architecture. Further, genetic associations identified in one or few strains may reflect only the penetrance of a given variant within a specific genetic background (Sittig et al. 2016). Comparing traits using large numbers of inbred strains allows us to resolve this confounding between causal mechanisms, but with significant effort and cost. Outbred populations randomize the genetic background to offer another alternative, but at the cost of large samples needed to test for differences between extreme trait values. Accurate trait predictions offer an approach to minimize these costs, enabling the sampling and analysis of extreme trait values without the need for large samples.

RATTACA presents a novel paradigm for sampling extreme trait values to facilitate experimental phenotyping and genetic correlations. This prediction-based framework in effect selects sets of individuals analogous to selected lines drawn from a single outbred population, eliminating any confounding between distinct causative effects underlying different traits. Here, we employed this framework on four traits ranging from modest (*h^2^* = 0.13) to high (*h^2^* = 0.47) narrow-sense heritability (Fig. 2B). In all cases, we successfully distinguished between samples with distinct mean trait values with high statistical significance (*P* = 1.09 x 10^-4^ −4.87 x 10^-35^). These successes suggest this framework is widely applicable. Given a sufficiently large sample on which to train models, G-BLUP prediction may reliably distinguish between phenotypically extreme samples even for low *h^2^* traits and modest genomic samples of tens of thousands of SNPs. As such, we suggest RATTACA is a robust and viable strategy for experimental sampling in HS rats, and an encouraging prospect for similar endeavors in other model systems. Further investigations testing different traits and samples, particularly in different taxa, will be needed to confirm the broader utility of this research paradigm.

### Model performance and validation

Heritability and sample size had the most obvious influence on our trait predictions. Model performance invariably improved with increasing *h^2^* and training population size (Fig. 3). These trends are consistent with a wide range of empirical and theoretical studies using multiple different prediction methods (Zhang et al. 2017; Liu, Xia, and Lan 2022; Wu et al. 2022). SNP sample size also covaried positively with model performance, but only across smaller genomic samples. Prediction accuracy increased for all traits with increasing SNP count until ∼10,000 SNPs, above which further SNP sampling had little to no apparent effect on model performance (Fig. 3B). Multiple prior studies have achieved modest to high prediction accuracy using reduced SNP samples (Li et al. 2018; Yoshida et al. 2018; Ober et al. 2012). Such cases likely reflect adequate sampling of causative SNPs affecting the phenotype, thus our asymptotic prediction accuracy suggests an upper limit to the number of loci underlying each of the four focal traits of this study. Increasing SNP counts above this limit did not increase variance in model performance (Fig. 3B), suggesting these additional SNPs accurately reflect realized relationships in our population. That is, because our G-BLUP models are equivalent to those based on an estimated genomic relationship matrix (Goddard 2009; Endelman 2011), these extra variants incur no penalty on model prediction accuracy, however they do increase the computation required for prediction.

We found no difference in prediction accuracy between models fit to randomly derived SNP samples vs. LD-pruned samples (Fig. 3C). Recombination rates in HS rats are modest (mean 0.66 cM MB^-1^) (Littrell et al. 2018), suggesting moderate long-range LD in this population (Munro et al. 2022); (Sanches et al., n.d.). As such, a given randomly sampled variant is moderately likely to be linked with a causative SNP for a given trait, meaning a relatively small random sample could adequately capture sufficient genetic variance underlying the phenotype to produce a good prediction (given heritability and population sample size constraints). This is in fact the pattern we found, with approximately ‘saturated’ prediction accuracy for models using random genomic samples at or above 10,000 SNPs (Fig. 3B). Below this threshold, prediction accuracy improved with increasing sampling of SNPs in LD with causal polymorphisms. Above this threshold, random sampling appears to capture approximately all causal variants, either directly or through linkage.

By capturing sufficient causal genetic variance, we can successfully distinguish between extreme trait samples using only a modest SNP count (Fig. 2B). This result is an encouraging sign that trait predictions can reliably be used for *a priori* experimental design without the need for computationally taxing analyses on large genomic datasets. Fortunately, this sampling paradigm also appears robust to a wide range of prediction accuracies. Our lowest-heritability trait (NRR, *h^2^* = 0.13) displayed consistently low prediction accuracy (max *r* < 0.2). Such model performance would likely be unreliable in other contexts requiring high confidence in individual-level predictions, such as genomic selection of genotypes for breeding or personalized medicine based on polygenic risk scores (Dudbridge 2013). However, RATTACA employs a more general sample-based selection paradigm in which individuals are selected for inclusion in an experimental sample based on their prediction rank, not pursued individually for their absolute predicted breeding value. As such, even with relatively low prediction accuracy RATTACA can successfully produce distinct experimental samples, as validated by significantly different observed trait means between selected samples (Fig. 2B). Traits with higher *h^2^* and for which we have more available experimental observations present the ideal usage cases for genetic prediction, however our analyses collectively suggest that RATTACA can be employed more broadly. As a general rule, we expect RATTACA predictions to reliably succeed for higher-heritability traits (*h^2^* > 0.2) with sample sizes greater than 300 individuals, and for lower-heritability traits (*h^2^* = 0.1 - 0.2) with samples greater than 600. The Center for Genetic Studies in Outbred Rats has data for >80 total traits meeting these criteria (Supplementary Material), which we offer as part of our prediction services to collaborators.

### RATTACA use cases: genetic correlation and more

RATTACA’s novel genetic prediction paradigm for experimental sampling opens a new avenue for identifying causal mechanisms shared between traits. Samples selected via RATTACA as having either high or low extreme predicted trait values are defined and differentiated by their genotypes, meaning individuals from one sample must be more closely related to each other than they are to the alternative sample (on average). Critically, because RATTACA samples are drawn from a single population, this differentiation cannot be attributed to confounding demographic factors that arise between separately bred laboratory strains. Therefore, if a pair of RATTACA samples is observed to differ for a second trait (not the trait used for prediction), this observed correlation is likely to have arisen due to a causative genetic architecture shared with the predicted trait. All simple trait correlations observed between RATTACA samples are thus valid candidates for genetic correlation.

Identifying candidate traits with a shared genetic basis is a simple process using RATTACA. A genetic correlation suggested by differentiation in a second trait means that only one trait need be measured (and one predicted) to infer shared causality. Between-sample differentiation in the measured trait can be assessed using a simple t-test between groups. A statistical difference between samples suggests a possible genetic correlation with the predicted trait used to define the samples. These tests need not adjust for relatedness with the likes of a genetic relationship matrix as our predictions are modeled on a surplus of SNPs (in addition to causal variants) that reflect population-wide ancestry, and because our sampling algorithm minimizes the number of sibships in a given sample. By controlling for these confounding factors in its sampling design, RATTACA makes it easy to identify candidate genetic correlations. Confirming these correlations will require concerted follow-up investigations beyond the scope of this paradigm.

One of RATTACA’s most compelling applications is its ability to help investigators circumvent undesired environmental effects on phenotypes of interest. For example, in some cases, measuring one trait can induce a change in another trait in a systematic way that is not due to genetic correlation. As an example, researchers pursuing the genetics of addiction may wish to compare genotypes with higher and lower propensities for addiction (addiction indices, e.g. (Carrette et al. 2022)). However, identifying samples with a given addiction index experimentally requires drug exposure, an environmental effect that may influence a second trait of interest (e.g., any number of behavioral or physiological correlates with drug exposure). RATTACA offers an opportunity to pursue correlations between traits without such experimental artifacts. For example, using genetic predictions of addiction index could enable comparisons between samples with extreme addiction propensities in which all individuals are naive to drug exposure, eliminating confounding environmental effects on the new trait of interest. RATTACA provides flexibility for further novel applications, such as when phenotypes are highly labor intensive and low-throughput to measure experimentally. In such cases, accurate trait predictions may be an easier, cheaper, preferred alternative for some investigators.

As a research service, RATTACA offers investigators rats and their respective trait predictions for use in an array of experimental contexts at low cost. This service is feasible given the established operation of the HS West colony which provides these rats. Maintaining a large outbred colony only to genotype a small subset of individuals for genotyping and prediction would be cost-prohibitive on its own. However, the large (more randomly selected) samples produced by HS West for various GWASs means RATTACA genotyping can proceed in bulk batches pooled with other projects in a cost-effective manner. Collectively, this laboratory population and sampling, genotyping, and prediction paradigm make RATTACA an economical and validated service that opens novel strategies for experimental sampling, genetic correlation, and more in HS rats. We outline this service and its early successes to highlight the utility of this novel framework in identifying connections between genotype and phenotype.

## Notes

### Competing Interest Statement

The authors have declared no competing interest.

