## Supplemental methods and tables for "RATTACA: Genetic predictions in Heterogeneous Stock rats offer a new tool for genetic correlation and experimental design"

### Genomic data preparation

The following detailed protocols are used in preparing RATTACA genotypes.

- DNA extraction: **dx.doi.org/10.17504/protocols.io.8epv59reng1b/v1**
- Library preparation: **dx.doi.org/10.17504/protocols.io.j8nlkkm85l5r/v1**
- Library normalization: **dx.doi.org/10.17504/protocols.io.261genw5dg47/v1**
- Library pooling and sequencing: **dx.doi.org/10.17504/protocols.io.yxmvmnw29g3p/v1**

**Table S1. Traits available for genetic prediction using RATTACA.** All traits have either reliable training sample size (N), narrow-sense heritability ($h^2$), or both required for reliably accurate prediction and sample selection.

| Behavioral Domain | Trait Category | Drug | Trait | Age (d) | N | $h^2$ |
|---|---|---|---|---|---|---|
| Addiction | Short access and Intermittent access to cocaine | Cocaine | Short access to cocaine, total infusions | 100 | 446 | 0.23 |
| Addiction | | Cocaine | Intermittent access, total infusions | 100 | 446 | 0.19 |
| Addiction | | Cocaine | Post-cocaine anxiety | 100 | 378 | 0.36 |
| Anxiety | | none | Baseline anxiety | 80 | 384 | 0.21 |
| Addiction | | Cocaine | Incentive sensitization - breakpoint | 100 | 446 | 0.10 |
| Locomotor | Locomotor | none | Total distance over 60 min | 45 | 1246 | 0.38 |
| Anxiety | | none | Duration in the center over 60 min | 45 | 1246 | 0.28 |
| Sensation seeking | Novelty seeking | none | Transitions to center zone frequency | 46 | 1245 | 0.21 |
| Sensation seeking | | none | Duration in center zone | 46 | 1245 | 0.25 |
| Sensation seeking | | none | Distance traveled in novel environment | 46 | 1245 | 0.29 |
| Sensation seeking | Novelty seeking | none | Transitions to novel environment zone | 60 | 1296 | 0.36 |
| Sensation seeking | | none | Distance traveled in novel environment | 60 | 1296 | 0.28 |
| Social Reinforcement | Social reinforcement | none | Duration in social zone | 47 | 1844 | 0.23 |
| Social Reinforcement | | none | Latency to enter social zone | 47 | 1328 | 0.14 |
| Anxiety, sensation seeking | Elevated plus maze | none | Duration in open arm | 48 | 1317 | 0.31 |
| Anxiety | | none | Duration in center | 48 | 1317 | 0.35 |
| Addiction | Socially acquired nicotine self administration | Nicotine | Total infusions | 55 | 1422 | 0.21 |
| Addiction | | Nicotine | Ratio between active and inactive licks | 55 | 1422 | 0.24 |
| Addiction | | Nicotine | Progressive ratio | 55 | 1422 | 0.16 |
| Addiction | | Nicotine | Reinstatement | 55 | 1422 | 0.10 |
| Addiction | Conditioned place preference to cocaine | Cocaine | Change in locomotor activity after cocaine | 140 - 204 | 1674 | 0.04 |
| Addiction | | Cocaine | Post-test time on conditioned side | 140 - 204 | 1674 | 0.04 |
| Learning | Conditioned reinforcement | none | Lever presses during conditioned reinforcement | 140 - 204 | 1628 | 0.22 |
| Learning | | none | Magasine responses during conditioned reinforcement | 140 - 204 | 1628 | 0.24 |

| | | | | | | |
|---|---|---|---|---|---|---|
| Learning | Conditioned reinforcement | none | Lever presses during conditioned reinforcement | 60 | 1583 | 0.24 |
| Learning | | none | Magasine responses during conditioned reinforcement | 60 | 1592 | 0.24 |
| Attribution of incentive salience to reward cues | Pavlovian conditioning | none | PavCA index score | 140 - 204 | 1645 | 0.20 |
| Attribution of incentive salience to reward cues | | none | PavCA latency score | 140 - 204 | 1645 | 0.22 |
| Attribution of incentive salience to reward cues | | none | PavCA probability difference | 140 - 204 | 1645 | 0.19 |
| Attribution of incentive salience to reward cues | | none | PavCA response bias | 140 - 204 | 1645 | 0.15 |
| Attribution of incentive salience to reward cues | Pavlovian conditioning | none | PavCA index score | 65 | 1583 | 0.23 |
| Attribution of incentive salience to reward cues | | none | PavCA latency score | 65 | 1591 | 0.22 |
| Attribution of incentive salience to reward cues | | none | PavCA probability difference | 65 | 1583 | 0.21 |
| Attribution of incentive salience to reward cues | | none | PavCA response bias | 65 | 1583 | 0.18 |
| Locomotor | Cocaine contextual conditioning | none | Locomotor distance baseline | 75 | 1245 | 0.23 |
| Addiction | | Cocaine | Locomotor distance after cocaine conditioning | 75 | 1336 | 0.24 |
| Addiction | Short access and Long access to heroin | Heroin | LgA: Long access total heroin intake | 105 - 130 | 862 | 0.22 |
| Addiction | | Heroin | Progressive ratio breakpoints | 130 | 862 | 0.13 |
| Addiction | | Heroin | Extinction bursts | 135 | 862 | 0.12 |
| Anxiety | | none | EPM duration in open arm baseline | 75-90 | 850 | 0.17 |
| Addiction | | Heroin | EPM duration in open arm after heroin | 135 | 850 | 0.08 |
| Locomotor | | none | OFT distance baseline | 75-90 | 858 | 0.15 |
| Addiction | | Heroin | OFT distance after heroin | 135 | 858 | 0.26 |
| Pain sensitivity | | none | Tail flick baseline | 75-90 | 847 | 0.19 |
| Analgesia | | Heroin | Tail flick after single heroin injection | 135 | 846 | 0.06 |
| Addiction | Short access and Long access to oxycodone | Oxycodone | LgA: Long access intake | | 530 | 0.12 |
| Addiction | | Oxycodone | ShA: Short access intake | | 308 | 0.12 |
| Addiction | | Oxycodone | Addiction index | | 530 | 0.02 |
| Pain sensitivity | | none | Tail flick baseline | | 528 | 0.17 |

| | | | | | | |
|---|---|---|---|---|---|---|
| Analgesia | | Oxycodone | Tail flick after oxycodone | | 528 | 0.10 |
| Pain sensitivity | | none | Von Frey baseline | | 527 | 0.18 |
| Analgesia | | Oxycodone | Von Frey after oxycodone | | 528 | 0.10 |
| Addiction | Short access and Long access to Cocaine | Cocaine | ShA: Short access intake last 2 days | | 790 | 0.08 |
| Addiction | | Cocaine | Addiction index | | 771 | 0.08 |
| Addiction | | Cocaine | Progressive ratio after ShA | | 787 | 0.11 |
| Addiction | | Cocaine | Irritability change after cocaine | | 372 | 0.12 |
| Locomotor | Cocaine avoidance and punishment resistance | none | Novelty-induced locomotion | | 743 | 0.31 |
| Shock escape | | Cocaine | Punishment resistance | | 982 | 0.30 |
| Addiction | | Cocaine | Progressive ratio | | 891 | 0.07 |
| Delay discounting | Delay discounting | none | K exponential curve fit | | 644 | 0.24 |
| Locomotor | Locomotor activity | none | Locomotor total distance | | 629 | 0.41 |
| Locomotor | | none | Locomotor rearing | | 629 | 0.26 |
| Locomotor | Locomotor activity | none | Locomotor total distance | | 1245 | 0.05 |
| Locomotor | | none | Locomotor rearing | | 1245 | 0.05 |
| Sensation seeking | Novelty seeking | none | Distance traveled in a novel environment | | 2509 | 0.27 |
| Sensation seeking | Light cue reactivity | none | Active snout-poke responses | | 2405 | 0.28 |
| Impulsivity, habituation | | none | Inactive snout-poke responses | | 2406 | 0.13 |
| Attention | Choice reaction time test | none | Mean reaction time | | 2453 | 0.20 |
| Attention | | none | Omissions | | 2454 | 0.22 |
| Impulsivity | | none | False alarms | | 2453 | 0.18 |
| Tibia length | Musculoskeletal | varies | Tibia length | 52 - 279 | 3516 | 0.48 |
| Muscle mass | | varies | Soleus mass | 52 - 279 | 3513 | 0.41 |
| Muscle mass | | varies | Extensor digitorum longus mass | 52 - 279 | 3510 | 0.52 |
| Muscle mass | | varies | Tibialis anterior mass | 52 - 279 | 3514 | 0.47 |
| Bone composition | Musculoskeletal | varies | Bone surface | | 900 | 0.05 |
| Bone composition | | varies | Bone density | | 900 | 0.05 |
| Bone composition | | varies | Bone cortical porosity | | 900 | 0.05 |
| Bone composition | | varies | Bone cortical thickness | | 900 | 0.05 |
| Cecal microbiome | Microbiome and metabolome | varies | multiple | | 1600 | ≤ 0.15 |
| Cecal metabolome | | varies | multiple | | 1000 | ≤ 0.15 |

**Table S2. Per-trait comparisons between model prediction accuracy using either randomly sampled SNPs or LD-pruned SNPs.** Estimated marginal means (EMMs) were estimated from a linear model of the form: performance ~ trait + prune method. EMMs and differences in EMMs are presented for each individual trait. The estimated difference, t-ratio and *P* value of the differences for the entire model suggest no statistical difference in model performance using either randomly sampled or LD-pruned SNP genotypes for trait prediction.

| Trait | Random EMM | LD Prune EMM | Trait Estimate (LD - Random) | Model Estimate | t-ratio | *P* value |
|---|---|---|---|---|---|---|
| Mass | 0.392 (0.386-0.398) | 0.394 (0.388-0.400) | 0.002 | 0.00255 | 0.932 | 0.352 |
| DNE | 0.300 (0.294-0.306) | 0.303 (0.297-0.309) | 0.003 | 0.00255 | 0.932 | 0.352 |
| VSSR | 0.284 (0.278-0.290) | 0.286 (0.280-0.292) | 0.002 | 0.00255 | 0.932 | 0.352 |
| NRR | 0.175 (0.169-0.181) | 0.178 (0.172-0.184) | 0.003 | 0.00255 | 0.932 | 0.352 |
